# Aperiodic EEG predicts variability of visual temporal processing

**DOI:** 10.1101/2023.12.05.570074

**Authors:** Michele Deodato, David Melcher

## Abstract

The human brain exhibits both oscillatory and aperiodic, or 1/f, activity. Although a large body of research has focused on the relationship between brain rhythms and sensory processes, aperiodic activity has often been dismissed as noise and functionally irrelevant. Prompted by recent findings linking aperiodic activity to the balance between neural excitation and inhibition, we investigated its effects on the temporal resolution of perception. We recorded EEG from participants during resting state and a task in which they detected the presence of two flashes separated by variable inter-stimulus intervals. Two-flash discrimination accuracy typically follows a sigmoid function whose steepness reflects perceptual variability or inconsistent integration/segregation of the stimuli. We found that individual differences in the steepness of the psychometric function correlated with EEG aperiodic exponents over posterior scalp sites. In other words, participants with higher levels of aperiodic activity (i.e., neural excitation) exhibited increased sensory noise, resulting in shallower psychometric curves. Our finding suggest that aperiodic EEG is linked to sensory integration processes usually attributed to the rhythmic inhibition of neural oscillations. Overall, this correspondence between aperiodic neural excitation and behavioral measures of sensory noise provides a more comprehensive explanation of the relationship between brain activity and sensory integration and represents an important extension to theories of how the brain samples sensory input over time.

## INTRODUCTION

Since Berger’s discovery of the alpha rhythm in EEG (1929), the study of brain oscillations and their potential roles in cognition has been a major focus of neuroscience research. Rhythmic patterns of brain activity are typically identified as peaks in the electroencephalography (EEG) power spectrum, with the alpha rhythm in the 8-12 Hz range being the most prominent. It has been proposed that alpha oscillations act as a sensory sampling mechanism, meaning that the duty cycle of the alpha rhythm (∼100 ms) corresponds to a perceptual snapshot or a temporal integration window (Michel et al., 2022; Ronconi et al., 2017; VanRullen & Koch, 2003). In support for this idea, Samaha and Postle, for example, found that the temporal threshold for discrimination of two fast consecutive flashes (i.e., two-flash fusion threshold) correlated with the speed of alpha oscillations (Samaha & Postle, 2015). Other studies have found that the period of alpha oscillations predicts the threshold for temporal order judgements (Chota et al., 2021; Kristofferson, 1967), visual temporal integration (Battaglini et al., 2020; Deodato & Melcher, 2023; Drewes et al., 2022; Gray & Emmanouil, 2020; Wutz et al., 2018), auditory temporal integration (Neuling et al., 2012), and multisensory temporal binding windows (Cecere et al., 2015; Migliorati et al., 2020; Ronconi et al., 2023; Venskus & Hughes, 2021). From a physiological standpoint, EEG alpha oscillations represent waxing and waning of inhibitory neural activity in the brain (Romei et al., 2008). Thus, sensory parsing could be achieved by regularly pulsed neural inhibition (Mathewson et al., 2011; Mazaheri & Jensen, 2010). However, recent null findings challenge the view that sensory integration may depend solely on oscillatory processes (Buergers & Noppeney, 2022; Keitel et al., 2022; Ruzzoli et al., 2019). Specifically, it has been suggested that internal noise could also affect the temporal organization of perception (Deodato & Melcher, 2023; Yarrow et al., 2022).

Apart from peaks of periodic activity, the EEG power spectrum exhibits a predominant aperiodic component, where power decreases as frequency increases following a power-law or 1/f pattern, suggesting the presence of irregular or non-rhythmic brain activity (B. J. He et al., 2010). Indeed, 1/f activity has been observed not only in M/EEG recordings, but also in Local Field Potentials and functional Magnetic Resonance Imaging (Donoghue et al., 2020; B. J. He, 2011, 2014; Manning et al., 2009; Miller et al., 2009; Ray & Maunsell, 2011).

Recent studies have shown that aperiodic brain activity can provide important insights into brain functioning (Churchill et al., 2016; B. J. He, 2011) and disease (Pani et al., 2022), contrary to the notion that it represents functionally irrelevant noise (Stumpf & Porter, 2012). Most notably, an increase of aperiodic activity, or a ‘flatter’ spectrum, has been consistently reported in older individuals compared to young ones. This has usually been interpreted as greater neural excitation, and E:I imbalance (Gao et al., 2017) or, from a computational perspective, as an increase in ‘neural noise’ and impaired information processing, given that the flat power spectrum is similar to that of white noise signals and predicts slower and less accurate cognitive performances (Donoghue et al., 2020; W. He et al., 2019; Ostlund et al., 2021; Voytek et al., 2015).

While previous studies have focused on oscillations, we challenged the long-standing assumption that aperiodic EEG activity is irrelevant for visual processing and investigated its functional significance for the temporal resolution of perception in the two-flash fusion task (see figure 1). Oscillations and aperiodic activity may represent two sides of neural communication, whose balance directly affects perception and, more generally, cognition (Voytek & Knight, 2015). Specifically, we reasoned that alpha oscillations’ pulsed inhibition mechanism could be altered by different levels of excitation/inhibition balance. Greater ‘neural noise’ could affect the rhythmic temporal organization of the visual flow by increasing the variability of neural processing latencies (Yarrow et al., 2022) or weakening oscillatory coupling (Voytek & Knight, 2015). Importantly, at the behavioral level, a noisier or unreliable sensory parsing/organization process should be observable in less reliable psychometric temporal thresholds or shallower psychometric curves (Deodato & Melcher, 2023).

**Figure 1.**
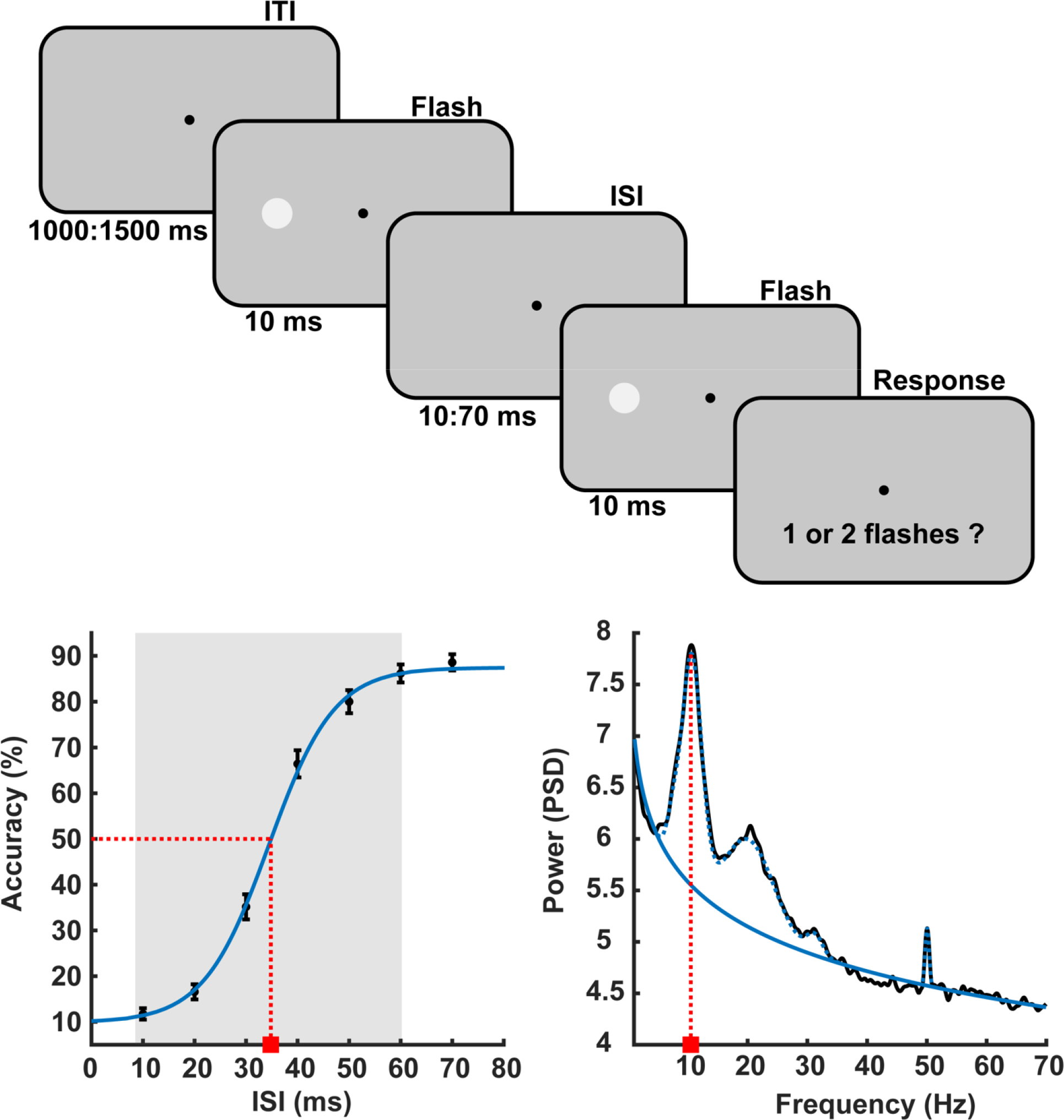
Experimental Design and Data modeling (top) Illustration of the two-flash fusion task (see methods). (bottom-left) Two-flash fusion accuracy is plotted against the inter-stimulus interval (ISI) (average across participants). The blue line indicates the psychometric function fit, the dotted red line indicates the threshold, and the shaded area indicates the function’ spread. Error bars indicate the standard error of the mean. (bottom-right) EEG spectrum of an example participant (channel Oz), where the solid blue line indicates the aperiodic fit, the dotted blue line indicates the periodic fit, and the dotted red line indicates the alpha frequency peak.

Overall, uncovering a role for the aperiodic component of EEG, in addition to oscillations, for temporal perception might provide an important extension to theories of rhythmic sensory sampling and a more comprehensive account of the relationship between brain activity and temporal integration/segregation, including a possible explanation of null findings.

## METHODS

### Participants

A total of 50 participants (26 females) between 18 and 50 years old (age mean = 21.62; std = 5.33) participated in the experiment and received compensation. Inclusion criteria were normal or corrected to normal vision and English fluency. Data were collected in accordance with the Declaration of Helsinki and the study protocol was approved by the local ethics committee (New York University Abu Dhabi IRB). Analysis of the same participants, in terms of oscillatory activity, has been previously reported (Deodato & Melcher, 2023).

### Experimental Design

At the beginning of the session there was a resting-stage EEG block in which participants were asked to let their mind freely wonder and relax for 5 minutes with closed eyes (EC condition) and then with open eyes (EO condition). Next, participants were familiarized with the two-flash task, in which they were instructed to fixate on the center of the screen and report if they perceived one or two flashes. The main experiment trials began with a central fixation on a gray background lasting for 1000-1500ms. This was followed by two consecutive flashes (∼20% contrast) presented on either the left or right side. Each flash lasted for 10ms and the interstimulus interval (ISI) was chosen from 7 equally spaced values between 10ms and 70ms (see figure 1). For each ISI, participants completed separately 10 training trials and 50 test trials. Trials were randomized and distributed in 2 blocks separated by a short break.

### Behavioral Data Analysis

The accuracy for each ISI was calculated by taking the average of the correct (i.e. two-flashes) responses. A psychometric logistic function was applied to the accuracy data for each individual participant and the fit of the model was determined using Cohen’s pseudo-R^2^ and a non-parametric likelihood-ratio test. Six participants were not included in the analysis since their lowest accuracy was above 30%, indicating that their responses were likely not due to temporal segregation (Boynton, 1972). One participant was excluded because the highest accuracy was 50%, and an additional five participants were removed due to poor model fit (<.90), leaving 38 participants for further analyses (see Deodato & Melcher, 2023).

### EEG Analysis

EEG data was recorded from 32 scalp active electrodes (Fp1, Fp2, F7, F3, Fz, F4, F8, FC5, FC1, FC2, FC6, T7, C3, Cz, C4, T8, TP9, CP5, CP1, CP2, CP6, TP10, P7, P3, Pz, P4, P8, PO9, O1, Oz, O2, PO10) using the BrainVision Recorder software. EEG was acquired with a sampling rate of 1000 Hz using the online reference at FCz. During pre-processing, an average of 0.5 bad channels per participant were detected and interpolated (range 0–6), Independent Component Analysis was used to identify and remove oculomotor artifacts. We avoided average reference because broadband noise in a noisy channel would be distributed across all channels and affect the shape of 1/f activity.

To compute the EEG power spectral density (PSD), the resting state data was divided in 2 second segments with 50% overlap and segments containing artifacts were rejected. The remaining segments were multiplied by a Hanning window, zero padded and Fourier transformed with a frequency resolution of 0.12 Hz, and converted to power. The aperiodic coefficient of the EEG was estimated with a linear fit of the log-log PSD spectrum after removal of any significant spectral peak (i.e., brain oscillations). The slope of the regression line in the log-log space corresponds to the negative exponent of the Lorentzian power spectral density function or scaling exponent. Alpha power was estimated for each electrode and participant after removing the aperiodic activity. Spectral parametrization was obtained with the “Fitting Oscillations and One Over F” (FOOOF) algorithm over the 1-70 Hz frequency range to avoid oscillations and spectral plateau crossing the fitting range borders (Donoghue et al., 2020; Gerster et al., 2022).

### Brain-Behavior Correlation

For each channel and resting state condition, the correlation between the EEG aperiodic exponent and the psychometric slope of the behavioral data was calculated across participants using the Pearson correlation coefficient. Notably, aperiodic exponents have been found to vary systematically with respect to age (Voytek et al., 2015) and alpha power (Muthukumaraswamy & Liley, 2018). Therefore, to control for the possible mediating effect of these variables, we conducted a mediation analysis (Baron & Kenny, 1986). First, aperiodic exponents and psychometric slopes were correlated with age and alpha power. Then, if a significant linear relationship was found, the regression between aperiodic exponents and psychometric slopes was recomputed while controlling for the effect of the intervening variable with multiple regression analysis (Baron & Kenny, 1986). Finally, to ensure that any result was not an artifact dependent on the parameters of the FOOOF spectral decomposition, we divided our sample in two groups (n=19) based on the steepness of the psychometric slope (i.e., “shallow” vs “steep” group) and compared their spectra frequency-wise with a cluster-corrected independent samples t-test.

Statistical significance of the results was established with non-parametric methods and cluster-based multiple comparisons correction (10000 permutations) (Maris & Oostenveld, 2007). Although we had *a priori* hypotheses in terms of the direction of the effects, all statistical tests were two-sided. Data analyses were implemented with MATLAB custom scripts, the Fieldtrip Toolbox (Oostenveld et al., 2011), and the FOOOF-mat toolbox (Donoghue et al., 2020).

## RESULTS

### Behavioral and spectral parametrization

In our sample (n=38), the temporal discrimination threshold of the two-flash fusion task (mean=35.2, std=6.57) was in line with previous research using similar paradigms to measure visual temporal resolution (Samaha & Postle, 2015). Although the fit of the sigmoid model was good and significant for all participants included in the data analysis (see figure 1 and methods), the shape of the function varied greatly across participants. This variability is best indicated by the slope (mean = .21, std = .102, range .044 - .5) or spread (mean = 42.82, std = 13.15, range 15.4 - 60) distributions (i.e., the range between the lower and upper asymptotes of the function), and could reflect individual differences in internal noise (Deodato & Melcher, 2023; Yarrow et al., 2022). Resting state EEG was recoded prior to the task and the “Fitting Oscillations and One Over F” (FOOOF: Donoghue et al., 2020) method was applied to achieve separation of periodic and aperiodic spectral components. This model provided excellent fits (mean R^2^ = .994). Values of the EEG aperiodic exponent (i.e., the spectral slope) ranged between 1.07 and 1.61 across channels (mean = 1.35; std = .15) and were greater over posterior scalp sites (see figure 2), corroborating previous reports of EEG aperiodic activity (Dehghani et al., 2010; Donoghue et al., 2020). Notably, the aperiodic exponent was significantly smaller across the whole scalp in the EO condition with respect to the EC (p<.05, two-tailed, cluster corrected) (see figure 2).

**Figure 2.**
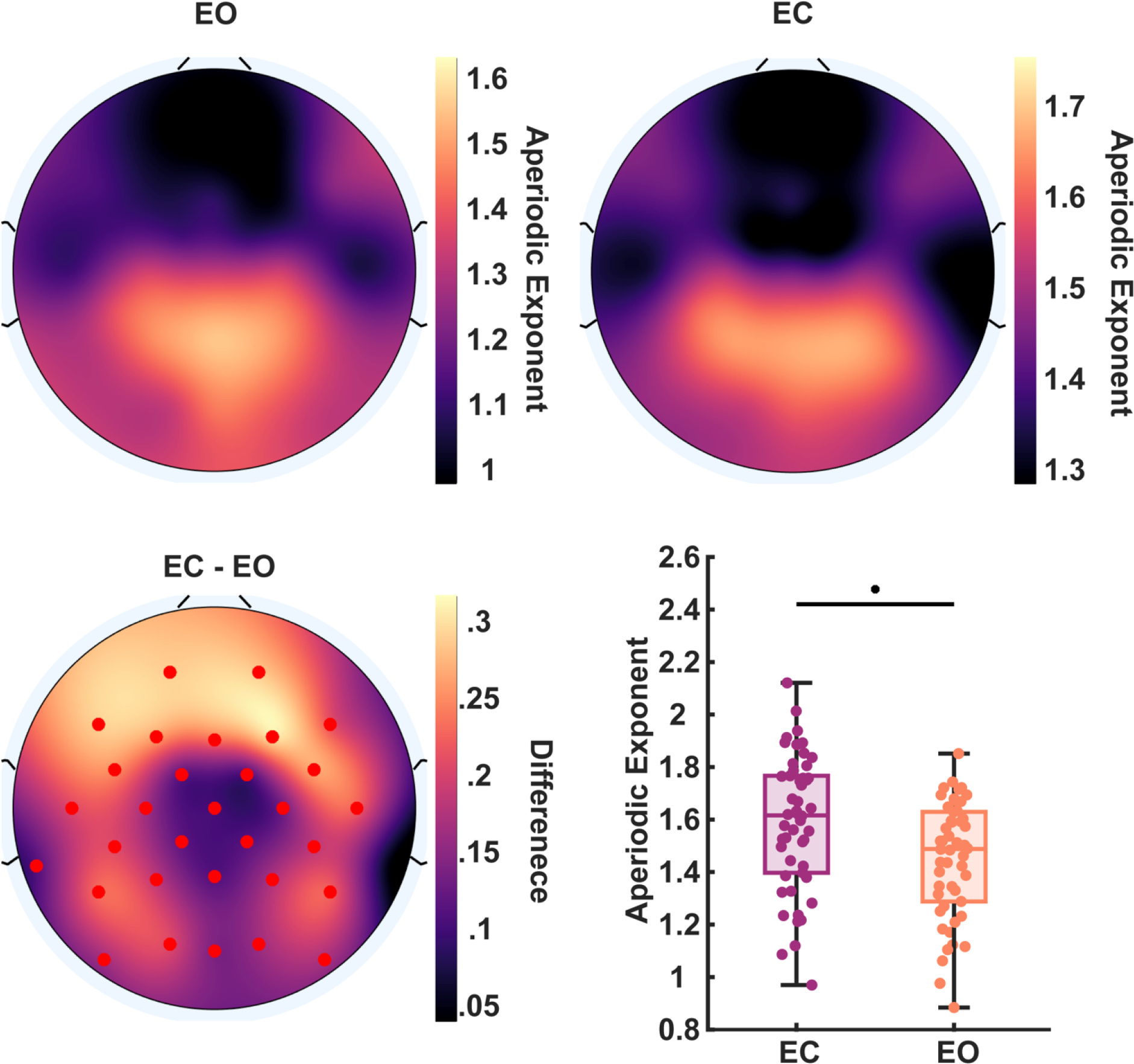
Aperiodic activity across the scalp. (top) Distribution of aperiodic exponents in the EO and EC resting state (averaged across participants) over the scalp. (bottom-left) Topographic map of the difference between the aperiodic exponents of EC and EO data, red markers indicate significant channels (p<.05). (bottom-right) Differences in the aperiodic exponent of EO vs EC resting state at channel Oz.

### Age- and alpha power-dependent changes in aperiodic EEG

As expected, we found a negative correlation between age and aperiodic exponents over posterior and central electrodes (see Figure 3). Older participants were characterized by a flatter power spectrum, replicating previous results concerning an age-dependent increase in neural noise (i.e., the neural noise hypothesis) (Voytek et al., 2015). Aperiodic exponents were also positively correlated with alpha power across participants, reflecting a trade-off between synchronous and asynchronous brain activity in the posterior region of the scalp (see Figure 3). While evidence for this relationship has already been reported both between and within subjects (Muthukumaraswamy & Liley, 2018), its underlying mechanism remains debated (Evertz et al., 2022; Muthukumaraswamy & Liley, 2018; Voytek & Knight, 2015).

**Figure 3.**
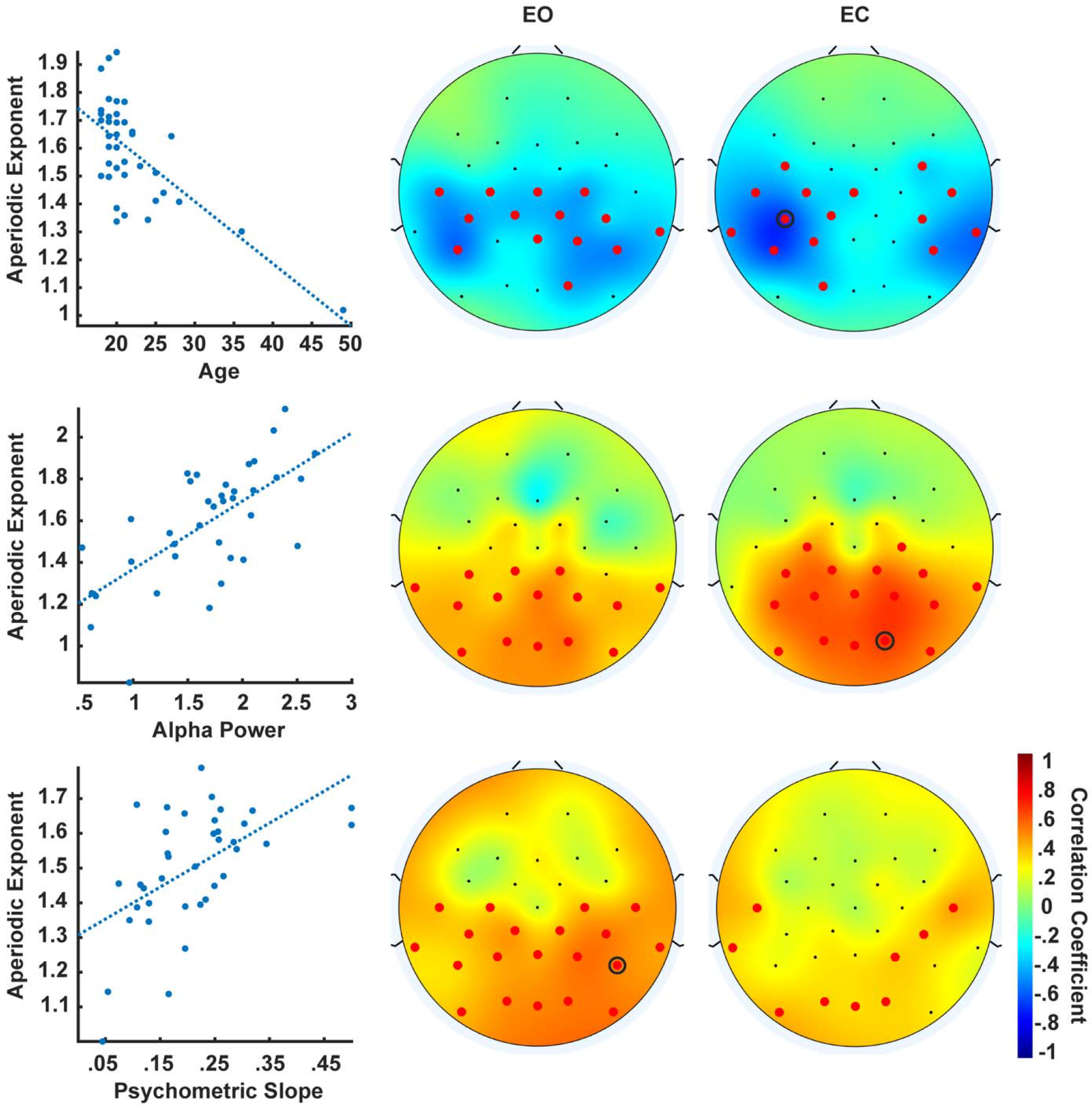
Correlation between two-flash fusion accuracy and resting state EEG. (top-row) Brain-behavior correlations between aperiodic exponents and age in the EO (middle) and EC (right) condition. The scatterplot shows the data and regression line (dotted line) for the channel with the greatest (absolute) correlation coefficient (black circle in the topographic maps). The topographic maps reflect the correlation coefficients, red markers indicate significant channels (p<.05). (middle and bottom rows) Same as top but for the correlations between aperiodic exponents and alpha power and two-flash fusion slopes.

### Aperiodic EEG correlates with two-flash fusion slope

The main correlation analysis revealed a positive association between the psychometric slope of the two-flash fusion task and the aperiodic coefficient of the resting state EEG (see Figure 3). Significant correlations emerged in the EC (mean r = .35; range = .32 - .44) and EO (mean r = .43; range = .32 - .55) conditions and were localized on the posterior region of the scalp. We found no relationship between psychometric slopes and participants age or alpha power (all p > .05), the results survived a mediation analysis controlling for the effect of these intervening variables. Thus, this correlation likely reflects a direct relationship between aperiodic activity and two-flash psychometric curve slopes. Additionally, a frequency-wise spectral comparison between participants with steep and shallow psychometric slopes revealed a posterior cluster of significant broadband differences above 30 Hz in both conditions, consistent with a change in the aperiodic exponent (see figure 4). These results indicates that participants with a flatter EEG spectrum had a shallower psychometric slope with respect to participants with greater aperiodic brain activity. In other words, greater aperiodic activity was associated with more variability in the two-flash integration/segregation task, consistent with an interpretation of neural noise.

**Figure 4.**
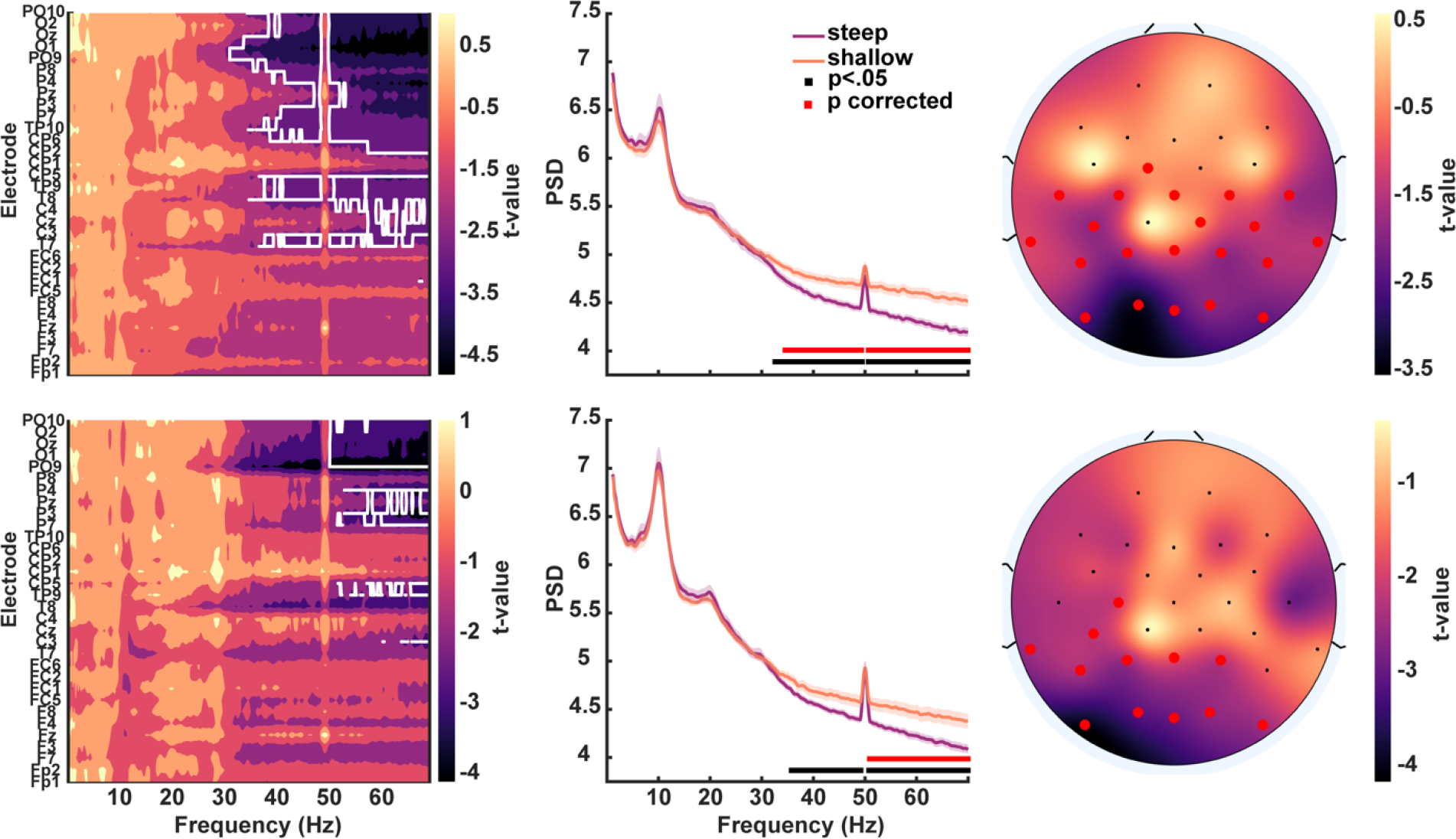
Spectral differences between participants with shallow and steep psychometric slopes. (left) Map of the t-values across frequencies and electrodes, the white outline indicates significant differences (p<.05) after multiple comparison correction. (middle) Comparison of the averaged EEG spectra of the two groups for channel Oz. (right) Topography of the t-values averaged between 30 and 70 Hz. Red markers indicate electrodes with significant differences. The top-row shows EO data, bottom-row shows EC data.

## DISCUSSION

We investigated the relationship between resting state EEG and performance in a visual temporal acuity task to shed light on the role of aperiodic neural activity in the temporal resolution of visual sensory processing. We found that discrimination of the stimuli depended on broadband arrhythmic brain activity during resting state. Specifically, greater aperiodic brain activity predicted shallower psychometric functions for two-flash fusion across participants, independently of age and oscillatory power.

These findings have important implications for understanding sensory processing. Current proposals suggest that discrimination of rapid visual events (e.g., two flashes) relies on a mechanism of sensory sampling implemented by neural oscillations in the alpha frequency range (∼10 Hz) (Samaha & Postle, 2015). Accordingly, using the same dataset we have previously shown that two-flash fusion thresholds correlate with the speed of alpha oscillations (Deodato & Melcher, 2023). Here, we hypothesized that aperiodic activity, which reflects greater neural excitation (Gao et al., 2017), could impair the rhythmic inhibition delivered by alpha oscillations, thus affecting the temporal organization of incoming sensory information (Mathewson et al., 2011; Mazaheri & Jensen, 2010). In other words, the neural noise and unspecific excitation indexed by 1/f neural activity might be related to temporal precision in neural processing, causing inconsistencies in the integration/segregation of the same stimulus across different trials.

Previous research has linked these inconsistencies with the steepness of the two-flash fusion slope (Deodato & Melcher, 2023; Gruzelier & Venables, 1974). Generally, a shallow slope indicates poor reliability of the psychometric threshold estimate because it implies a larger interval of mixed responses and greater inconsistency in perceiving the same stimuli. This aligns with evidence that participants with shallow slopes do not exhibit a clear relationship between psychometric thresholds and neural oscillations (Deodato & Melcher, 2023) and are generally less susceptible to the effect of oscillatory phase on perception (Ten Oever et al., 2020). Conversely, a steep psychometric slope reflects consistent integration/segregation responses for each inter-stimulus interval. Notably, it has been suggested that the steepness of the psychometric function directly reflects the quality of sensory processing. This internal noise hypothesis (Schneider et al., 1989) posits that greater neural variability would impair consistent stimulus processing and cause shallower psychometric slopes in detection tasks (Allen & Nelles, 1996; Buss et al., 2006, 2009; Sobon et al., 2019; Vilidaite & Baker, 2017). In a similar vein, Yarrow and colleagues showed that the steepness of psychometric functions describing multisensory temporal integration is associated with the variability of modality-specific neural processing latencies (Yarrow et al., 2022).

The EEG spectral slope (i.e., the aperiodic exponent) has been recently interpreted as a measure of neural noise and information processing due to signal processing considerations as well as empirical findings. Specifically, a flatter frequency spectrum indicates higher levels of white noise and reduced scale-free activity, both of which are detrimental to the quality of neural signals and communication (Miller et al., 2009; Voytek et al., 2015; Voytek & Knight, 2015).

Furthermore, flatter EEG spectra have been associated with impaired neuronal communication in clinical conditions (Pani et al., 2022; Voytek & Knight, 2015) and aging, supporting the neural noise hypothesis (Donoghue et al., 2020; Voytek et al., 2015; Waschke et al., 2017). Here, we propose that aperiodic activity and E:I balance, rather than being an unspecific index of neural noise, could be intrinsically related to oscillatory activity and the temporal organization of perception and, more generally, cognition. A similar proposal has been advanced by the Dynamic Network Communication framework: flattened power spectral slopes could result from increased background rates of neural firing caused by weak temporal coupling between neural spikes and the phase of oscillatory activity (Voytek & Knight, 2015). This explanation aligns with the observed trade-off between aperiodic activity and alpha power and further supports the finding that the desynchronized firing patterns commonly observed in flatter EEG spectra are an indicator of the balance between synaptic excitation and inhibition (Gao et al., 2017).

Furthermore, recent research suggested that individual differences in neural variability and aperiodic activity are reflected in the quality and speed of neural processing (Garrett et al., 2013). Ouyang and colleagues, for example, found that the EEG power law exponent predicted individual differences in cognitive processing speed in an object recognition task (Ouyang et al., 2020). Additionally, it has been reported that aperiodic EEG mediated age-related differences in cognition, but only in a task requiring speeded processing (Pathania et al., 2022). On a related note, scale-free properties of EEG have been found to mirror those of timing estimation errors in rhythmic tapping (Smit et al., 2013) and of detection of low-intensity flashes and sounds (Palva et al., 2013; Waschke et al., 2021), supporting a link with variability in the consistency and temporal pace of neural processing. This variability might have clinical implications in both general and visual processing-related conditions. For example, Turri and colleagues recently reported that individuals affected by developmental dyslexia, a condition characterized by abnormal spatio-temporal organization of the visual flow, display flatter EEG spectral slopes compared to controls (Turri et al., 2023).

Finally, we found significant differences between aperiodic activity of the EO and EC conditions (see figure 2), as well as qualitatively different scalp topographies for their brain-behavior correlations (see figure 3). The wide-spread association between aperiodic EEG and two-flash fusion accuracy in the EO, rather than EC, condition could be explained in light of the similarity of the EO condition to the task condition. Similarly, while we cannot rule out the influence of ocular artifacts on aperiodic differences between EO and EC, these state-related changes could be a physiological excitatory response to sensory processing (El Boustani et al., 2009) and arousal (Lendner et al., 2020). However, the finding that resting state in both conditions correlated with subsequent task performance suggests that aperiodic EEG exponents are a relatively stable trait across individuals. Indeed, scale-free properties of individuals’ brain activity have shown consistent test-retest variability and a link to the underlying genetics of participants (Linkenkaer-Hansen et al., 2007; Nikulin & Brismar, 2004), and have recently been considered a subject-specific signature that is precise enough to discriminate single individuals in large EEG datasets (Demuru & Fraschini, 2020).

In conclusion, the present study offers a clear link between individual differences of resting state aperiodic EEG (or neural E:I balance) and performance in a visual temporal discrimination task. We argue that heightened neural excitation could alter the rhythmic inhibition delivered by neural oscillations, causing inconsistent temporal segregation/integration of the visual stream. Overall, these results contribute to a growing literature highlighting the relevance of brain-related aperiodic activity in human cognition and behavior. As such, aperiodic activity should not be dismissed as nuisance noise, but considered as an important measure of *neural* noise and functional variability.

## Author Contribution

Michele Deodato: Conceptualization; Data curation; Formal analysis; Investigation; Methodology; Software; Validation; Visualization; Writing—Original draft; Writing—Review & editing.

David Melcher: Conceptualization; Funding acquisition; Methodology; Validation; Writing— Review & editing; Project administration; Resources; Supervision;

## Data Availability Statement

Data are publicly available in the Open Science Framework (osf.io/acb9n).

